# An evaluation of resonant scanning as a high-speed imaging technique for two-photon imaging of cortical vasculature

**DOI:** 10.1101/2021.11.11.468253

**Authors:** Annie Zhou, Shaun A. Engelmann, Samuel A. Mihelic, Alankrit Tomar, Ahmed M. Hassan, Andrew K. Dunn

**Affiliations:** Department of Biomedical Engineering, The University of Texas at Austin, 107 W. Dean Keeton C0800, Austin, TX 78712, USA

## Abstract

We demonstrate a simple, low-cost two-photon microscope design with both galvo-galvo and resonant-galvo scanning capabilities. We quantify and compare the signal-to-noise ratios and imaging speeds of the galvo-galvo and resonant-galvo scanning modes when used for murine neurovascular imaging. The two scanning modes perform as expected under shot-noise limited detection and are found to achieve comparable signal-to-noise ratios. Resonant-galvo scanning is capable of reaching desired signal-to-noise ratios using less acquisition time when higher excitation power can be used. Given equal excitation power and total pixel dwell time between the two methods, galvo-galvo scanning outperforms resonant-galvo scanning in image quality when detection deviates from being shot-noise limited.

## 1. Introduction

Two-photon fluorescence microscopy (2PM) is widely used in neuroscience since it enables non-invasive *in vivo* imaging with high lateral resolution on the order of microns and at depths exceeding 1 mm in brain tissue [1]. With traditional 2PM, imaging is completed by scanning a focused, ultrafast laser beam across a sample and measuring the resulting fluorescent signal. Like other point-scanning techniques, however, the temporal resolution of traditional 2PM is limited and collection of three-dimensional volumetric images can be time consuming.

Existing video-rate point scanning techniques include the use of resonant galvanometers, acousto-optic beam deflectors (AODs), digital mirror programming arrays (DMDs), and polygon scanning mirrors [2–5]. The use of these methods requires a compromise in microscope performance in addition to an increase the cost and complexity of the system compared to scanning with slower galvanometer mirrors. Using AODs is challenging due to its dispersive properties, which lead to limitations in the axial range. Other common challenges include minimizing spherical aberrations, which increase with axial range, and compensating for temporal pulse broadening [4,5]. One challenge with using DMDs is that the illuminating laser beam must be large enough to cover the entire surface of the DMD. Other challenges lie in optimizing for light throughput, axial resolution, offset rejection, and photobleaching, all of which are tradeoffs for one another, and characterization requires testing of numerous scanning and illumination patterns [2,3]. Polygon scanning mirrors requires a more complex instrument design and requires the complicated, optical correction of pyramidal errors, variations in reflectivity and angle that change with the axis of rotation [5]. Several alternatives to point scanning, such as SCAPE [6,7], reverberation 2PM [8], Bessel 2PM [9], FACED [10], rescanning using a scan multiplier unit [11], and two-photon light sheet microscopy [12], have been developed to increase temporal resolution of volumetric imaging. Compared to video-rate point scanning techniques, however, these methods come with even higher material cost and complexity.

Because of these limitations, resonant scanning remains the most common method for high-speed point scanning microscopy and advances in data acquisition hardware have accelerated its adoption. Field programmable gate array (FPGA) modules and open-source software for controlling this hardware, primarily ScanImage, have simplified high speed acquisition and correction for the sinusoidal scan speed of the resonant galvanometer [13]. High speed acquisition, however, significantly decreases pixel dwell times and necessitates higher excitation powers or frame averaging to achieve image quality that is comparable to slower, non-resonant scanning.

In this paper, we present a design for a two-photon microscope that includes a resonant galvanometer and a non-resonant galvanometer pair and experimentally compare the image signal to noise for *in vivo* vascular imaging under both types of scanning. The design for this custom-built resonant-galvo-galvo (RGG) two-photon microscope enables seamless switching between the two scanning methods and is straightforward to implement. We compare acquisition speeds between the two scanning methods for collecting images with the same signal-to-noise ratios at up to 750 μm depths in a mouse cerebral cortex. We also examine each method’s performance when imaging with identical excitation power and total sample exposure time. We demonstrate the ability to collect large field of view (FOV) images with up to fourfold speedup using resonant scanning, with image quality suitable for the processing and vectorization of the vasculature.

## 2. Materials and methods

### 2.1 Two-photon microscope design

The excitation source and upright microscope used for imaging were both custom-built. The source is an ytterbium fiber amplifier seeded by a low power commercial oscillator (Origami-10, OneFive GmbH, 100 mW, 80 MHz) that is amplified by a custom fiber amplifier resulting in an output beam of 1050 nm wavelength, 120 fs pulse width, 80 MHz repetition rate, and up to 6 W of average power [14,15]. The microscope, shown in Fig. 1, consists of an 8-kHz resonant galvanometer scanning mirror (CRS 8 kHz, Cambridge Technology) conjugated to an *xy*-galvanometer scanning mirror pair (GVS012, Thorlabs) using two identical scan lenses (*f*=50 mm, SL50-2P2, Thorlabs). The resonant scanner sits in a custom-designed, aluminum housing (Supplement 1). To avoid image distortion during resonant scanning (Fig. 2(c,e)), the resonant scanner is aligned using a tip, tilt, and rotation stage (TTR001, Thorlabs) to center the resonant scan along the axis of rotation of the first mirror in the galvo mirror pair and produce a square scan (Fig. 2(d,f)). A third scan lens (*f=*50 mm, SL50-2P2, Thorlabs) coupled with a Plössl tube lens (*f=*200mm, 2 × AC508-400-C, Thorlabs) expand the beam to fill the back aperture of the microscope objective (20x, 1.0 NA, XLUMPLFLN 20XW, Olympus). Heavy water was used for the immersion medium. Power is adjusted with an electro-optic modulator (EOM) before entering the microscope and did not exceed 170 mW at the sample to avoid tissue damage [16]. Fluorescence is epi-collected and passed through a 609/181 bandpass filter (FF01-609/181-25, Semrock) before reaching the photomultiplier tube (PMT, H10770PB-40, Hamamatsu Photonics). The PMT output is amplified with a variable gain transimpedance amplifier (DHPCA-100, FEMTO) before being passed through National Instruments hardware: digitizer, FPGA, and DAQ. Galvanometers, EOM, and image acquisition were controlled with ScanImage software (Vidrio Technologies) [13].

**Fig. 1.**
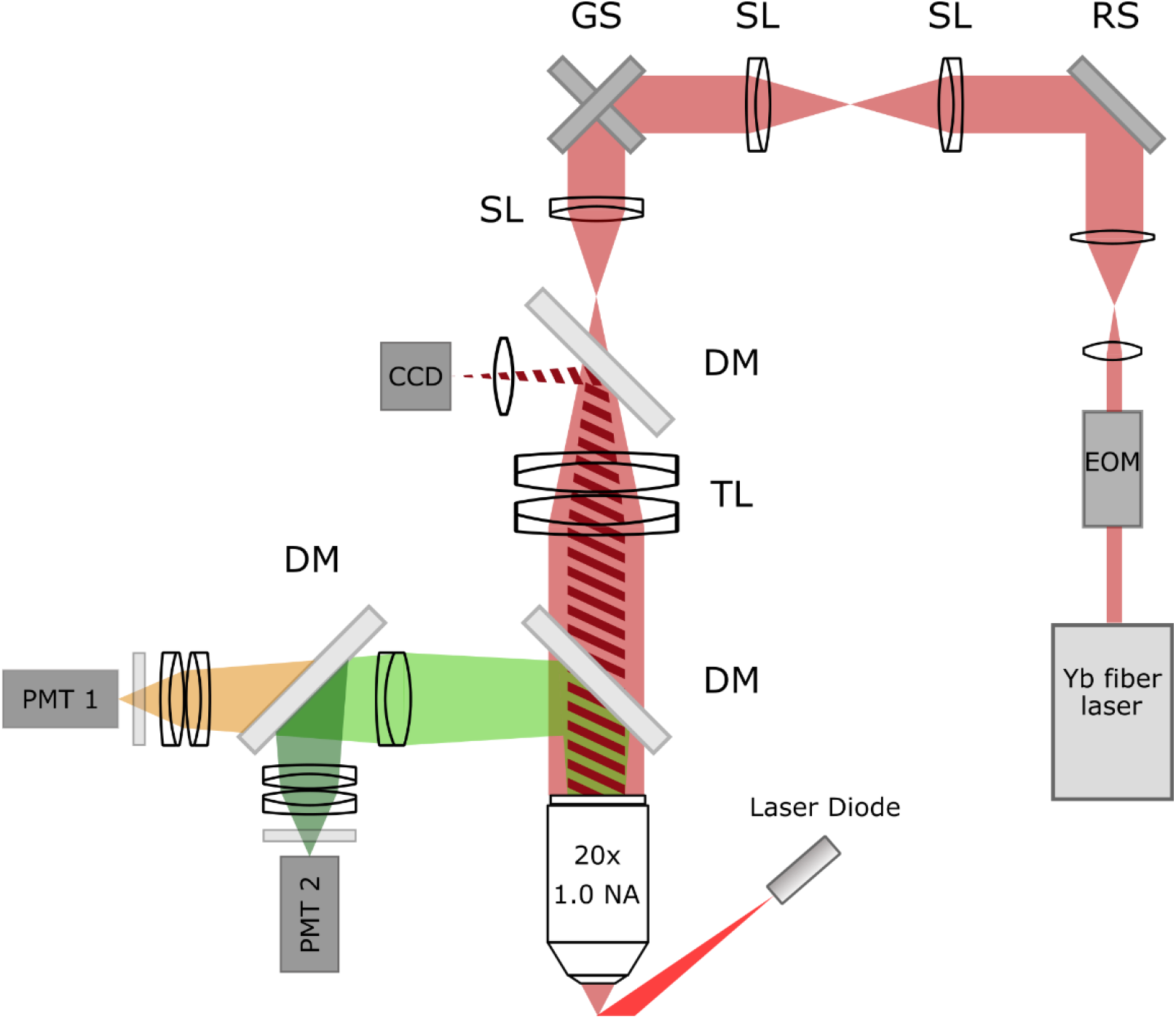
Resonant-galvo-galvo microscope schematic. EOM: elctro-optic modulator. RS: resonant scanner (on a tip, tilt, and rotation stage). SL: scan lens (*f*=50 mm). GS: galvanometer scanners. DM: dichroic mirror. TL: tube lens (*f*=200mm). PMT: photomultiplier tube. Excitation sources: Yb fiber laser (λ=1050 nm) for two-photon imaging, laser diode (λ=820 nm) for speckle imaging.

**Fig. 2.**
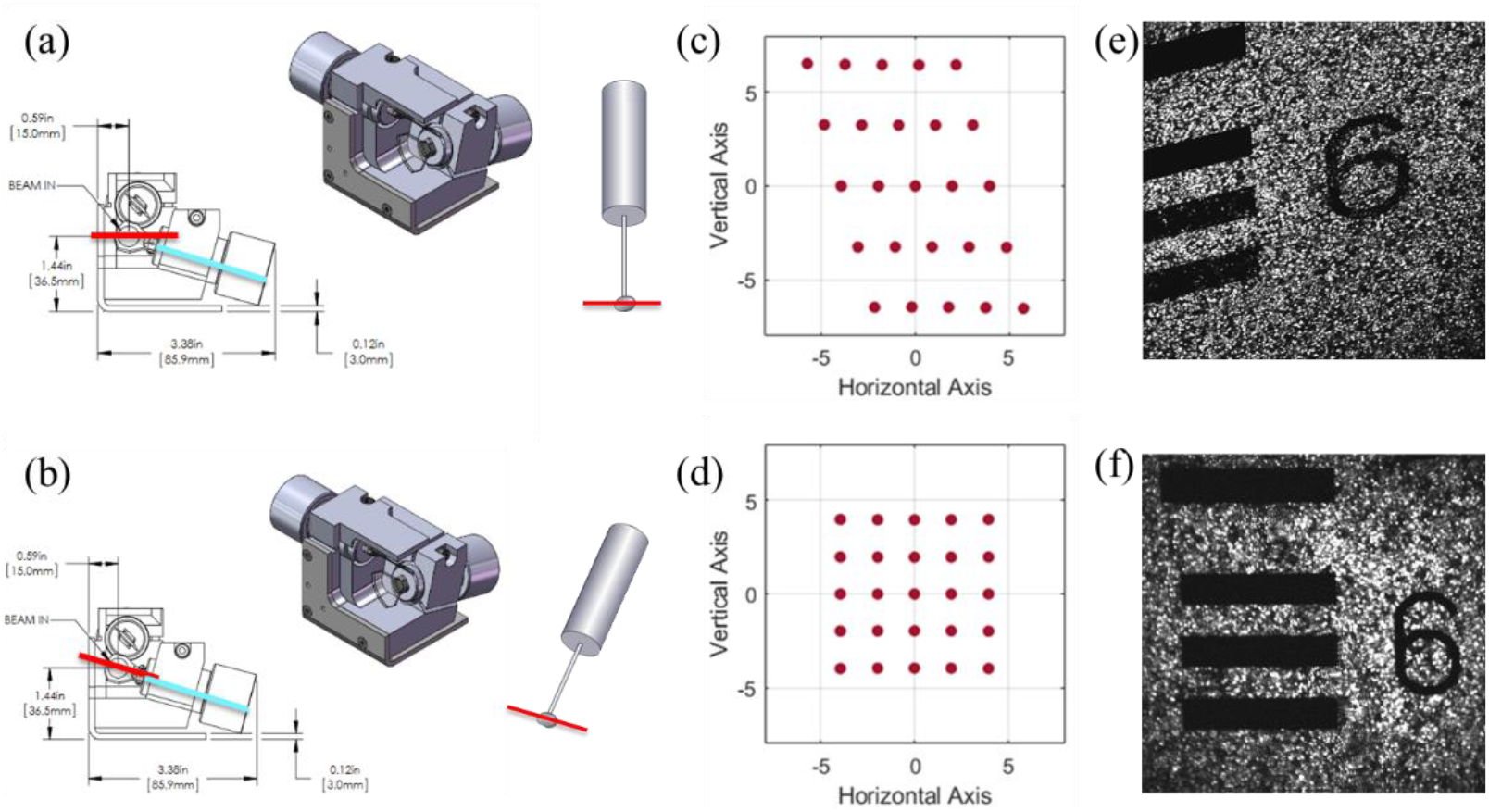
CAD drawing of GVS012 scanning galvanometer pair (courtesy of Thorlabs, Inc.) and schematic of resonant galvanometer with outgoing beam scan direction and angle from resonant galvanometer shown as a red line for comparison against the tilt angle of first galvanometer mirror shown in blue for (a) resonant galvanometer alignment with axis of rotation perpendicular to breadboard and (b) resonant galvanometer alignment with 15° tilt to align axis of rotation perpendicular to that of first galvanometer mirror. Resulting scans at the imaging plane for each configuration are shown as the output of a Zemax model for scan angles between -10 and 10 degrees in (c) and (d), and with sample images of a fluorescent USAF target in (e) and (f), for configurations (a) and (b) respectively.

The two-photon microscope design includes a laser speckle contrast imaging (LSCI) system that shares the objective used for two-photo imaging. With this setup, widefield LSCI images of blood flow and cortical surface vasculature can be acquired prior to two-photon imaging and used to determine ideal two-photon imaging locations [17–20]. Using the LSCI image to find regions further reduces the amount of time and laser exposure needed during the two-photon imaging session. To collect the LSCI image, the cranial window is illuminated using a laser diode (λ = 820 nm). The reflected light is passed through the objective and tube lens and is redirected with a dichroic mirror (FF875-Di01-25×36, Semrock) through a biconvex lens (*f*=100 nm) before reaching the CCD camera (acA2040-90umNIR, Basler AG).

### 2.2 Animal Preparation

Cranial window implants were prepared in C57 mice with dura intact, as previously described [20]. During imaging, mice were anesthetized with isoflurane and body temperature was maintained at 37.5 °C. Fluorescent labeling of blood plasma was completed with dextran conjugated Texas Red (70kDa, D1830, Thermo Fisher) dissolved in saline (5% w/v), administered intravenously via retro-orbital injection (0.1 mL). All animal protocols were approved by The University of Texas at Austin Institutional Animal Care and Use Committee.

### 2.3 Image Analysis

Images were rendered with Fiji ImageJ [21] and ParaView [22]. Quantitative image analysis was performed using MATLAB. Signal-to-noise ratios (SNRs) were determined for line profiles of 5-pixel width and 48-pixel length (approx. 7 μm × 65 μm) across representative vessels. SNR is quantified by (*µ*_*sig*_ − *µ*_*bg*_) / *σ*_*bg*_ where *μ*_*sig*_ is the mean signal intensity and *μ*_*bg*_, *σ*_*bg*_ are the mean and standard deviation of the background intensity, respectively. In SNR analysis, the acquisition time represents the total time spent scanning minus the total frame flyback time.

Image vectorization was performed using Segmentation-Less Automated Vascular Vectorization (SLAVV) software [23]. The program extracts vector sets representing the vascular network and calculates a variety of network statistics. The vectorized image can be displayed and examined using MATLAB or Blender add-on *VessMorphoVis* [24].

## 3. Experimental results

### 3.1 Alignment of resonant galvanometer

To avoid image distortion at the sample plane, the resonant galvanometer was aligned using a tip, tilt, and rotation stage. The appropriate tilt angle was determined from the CAD drawing of the galvanometer pair, which showed that the first galvanometer mirror in the path sits at a 15° angle from the horizontal plane (Fig. 2(a, b)). Analysis was performed in Zemax to confirm that a 15° roll angle would create a square image (Fig. 2(d)) and that orthogonal placement of the resonant galvanometer would create a distorted image (Fig. 2(c)). Images of a fluorescent US Air Force target taken before (Fig. 2(e)) and after (Fig. 2(f)) the tilt correction were consistent with Zemax analysis.

### 3.2 In vivo two-photon microscopy imaging

Two-photon microscopy images of cortical vasculature were collected using two different scanning methods for image quality comparison. Resonant-galvo (RG) scanning was performed with the resonant galvanometer scanning in the horizontal direction and the *x* galvo mirror of the *xy*-galvanometer pair scanning in the vertical direction. Galvo-galvo (GG) scanning was performed with the *x* and *y* galvanometer mirrors in the *xy*-galvanometer pair scanning in the horizontal and vertical directions, respectively, and the resonant galvo parked at its zero position. Excitation powers for each method were selected at each depth to be the power required to maximize the illumination of the vasculature without saturation, while minimizing the creation of out-of-plane fluorescence when viewing the image at 1 frame average for GG scanning and 20 frame average for RG scanning. To avoid tissue damage, power at the tissue surface was kept below 100 mW for GG scanning and below 170 mW for RG scanning [16].

Figure 3 demonstrates an image stack collected using RG scanning. An imaging depth of 750 μm was accomplished with 700 μm by 700 μm images collected at 3 μm depth increments, with optimized frame averaging determined by Figure 4 to yield a SNR of at least 35 (15 frames from 0 to 250 μm, 40 frames from 250 to 500 μm, 100 frames from 500 to 700 μm, and 360 frames from 700 to 750 μm). With a frame rate of 27 Hz, this stack was acquired in 14 minutes.

**Fig. 3.**
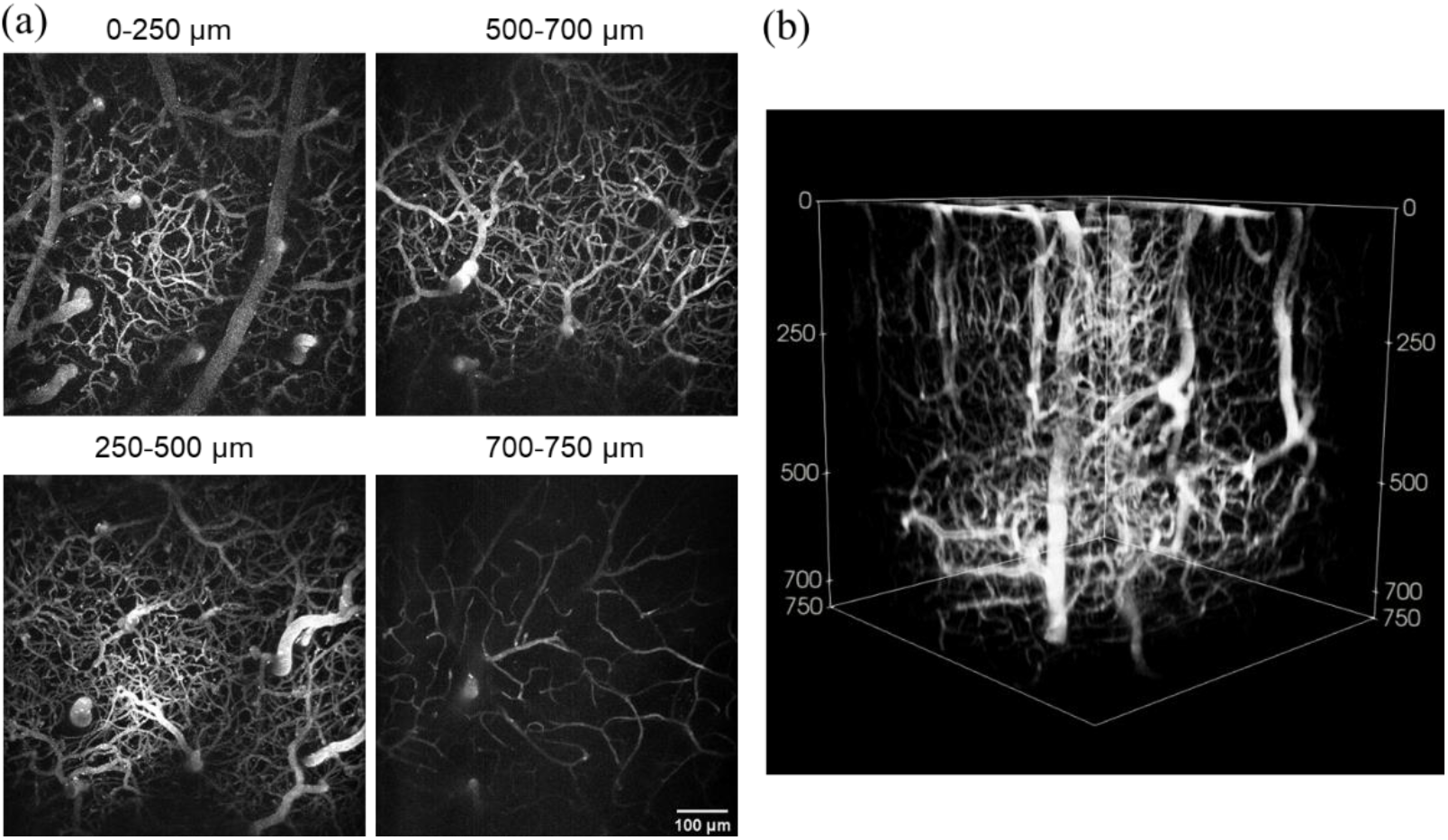
*In vivo* two-photon microscopy images of cortical vasculature taken with resonant-galvo scanning. (a) x-y intensity projections showing frame averages of 15 frames from 0 to 250 μm, 40 frames from 250 to 500 μm, 100 frames from 500 to 700 μm, and 360 frames from 700 to 750 μm. (b) Three-dimensional reconstruction of the stack from which the projections in (a) were taken.

**Fig. 4.**
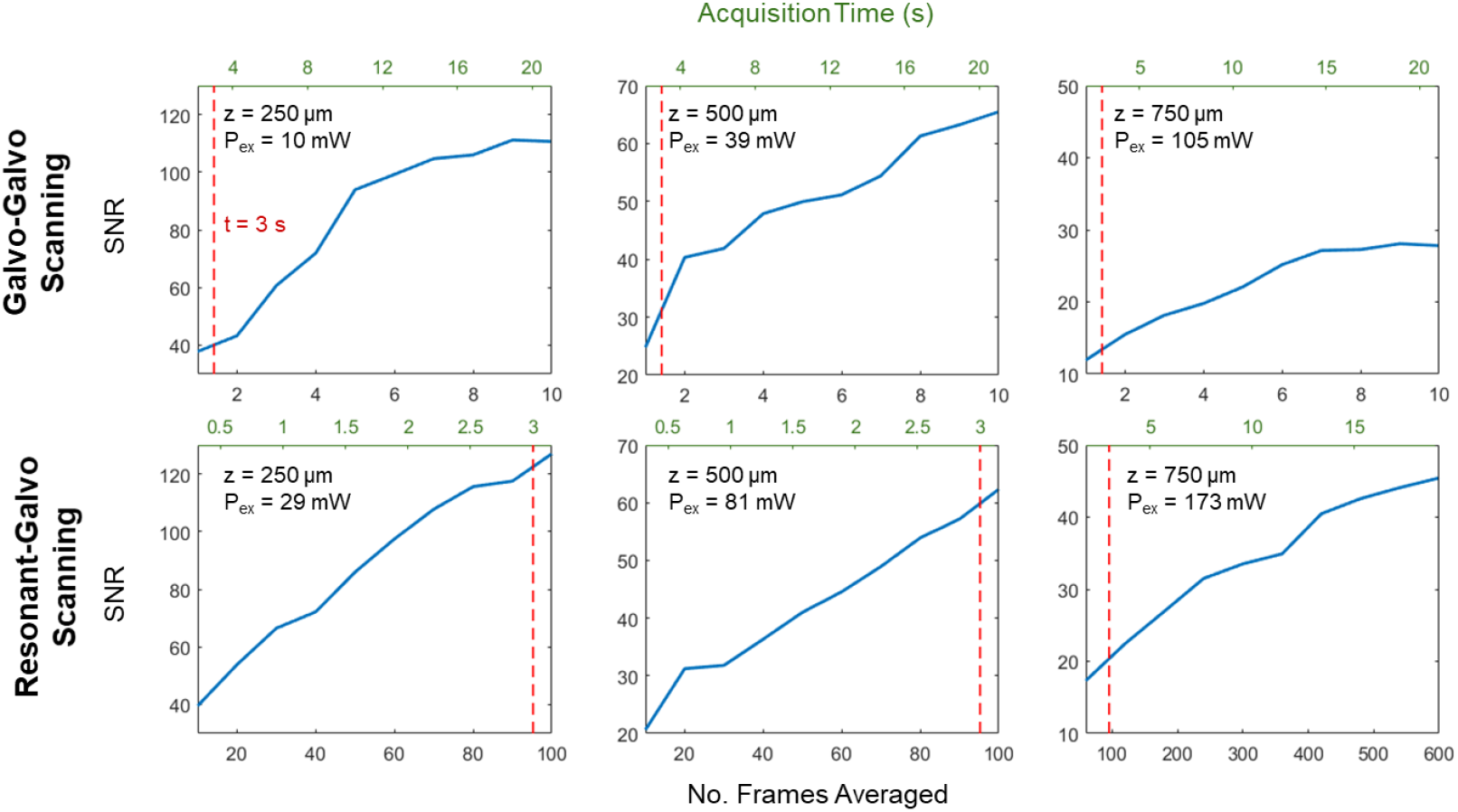
Comparison of SNR against number of frames averaged and corresponding acquisition times for resonant-galvo and galvo-galvo scanning methods. Each SNR value plotted is the average SNR across three representative vessels chosen at each depth *z*, (250, 500, and 750 μm), for which line profiles were analyzed at each level of frame averaging. *P*_*ex*_ is the excitation power at the sample for each acquisition. The frame rates were 0.48 Hz for GG scanning and 27 Hz for RG scanning.

To compare RG and GG scanning under typical *in vivo* imaging conditions, we calculated the SNR values for different levels of frame averaging for each scanning method as a measure of image quality per acquisition time. The average powers were greater for RG than for GG scanning in these measurements and are listed in Figure 4. Each SNR plot shown in Figure 4 is the average of SNR values across three representative vessels chosen at each depth (250, 500, and 750 μm), for which line profiles were analyzed at each level of frame averaging. Figure 5(a) demonstrates line profiles shown in yellow across of one of the three vessels chosen at each depth, with frame averaging chosen to give an SNR of approximately 35. Figure 5(b) plots the line profile results at 750 μm depth for various levels of frame averaging that are equivalent in scan time across the two scanning methods. Samples of resulting images from different levels of frame averaging for GG and RG scanning are shown in Figure 5(c), for scan times of 3, 13.5, and 22.5 s at 750 μm depth.

**Fig. 5.**
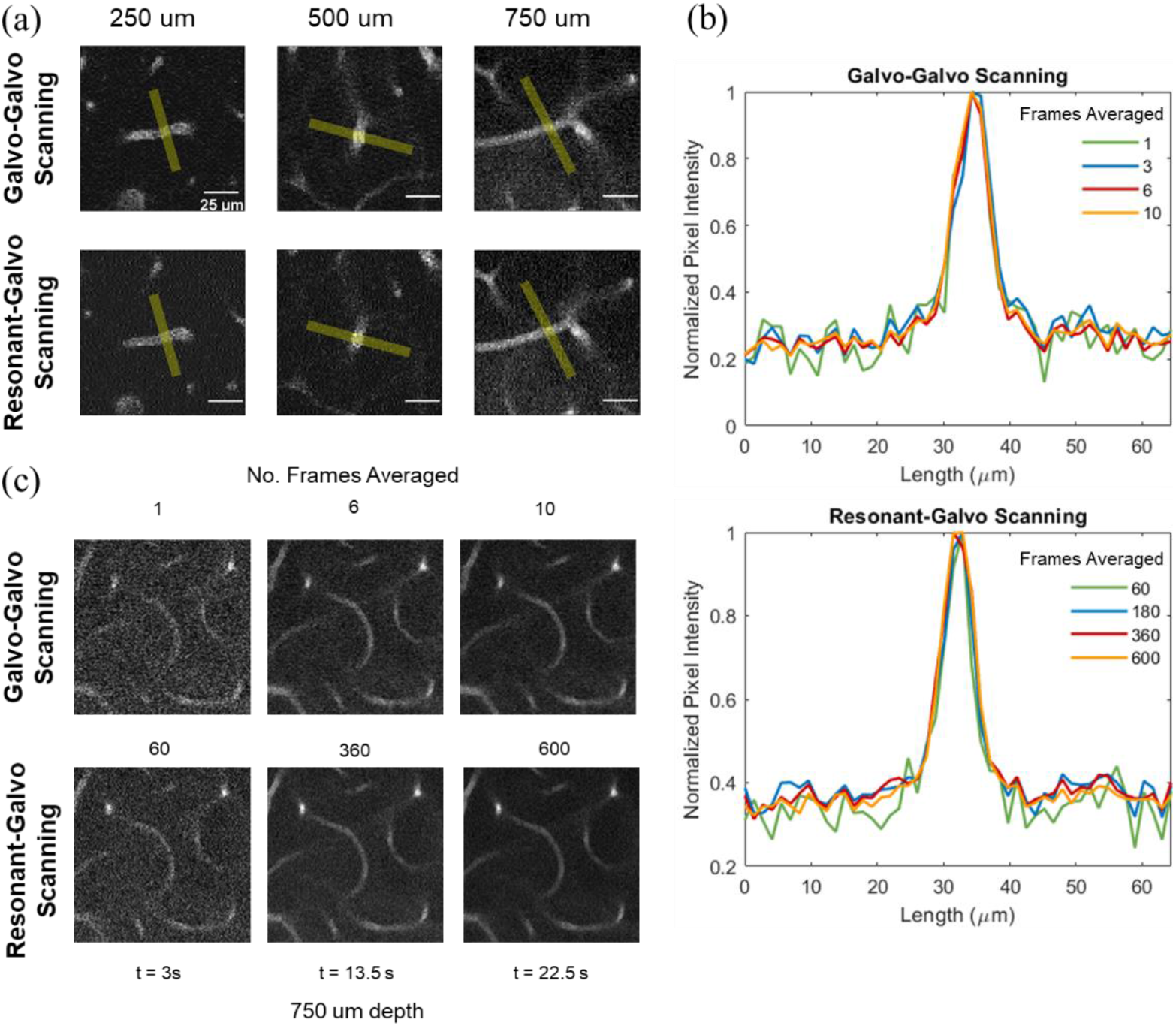
(a) Sample vessels and line profiles shown in yellow (approx. 7 μm × 65 μm) at each depth for RG and GG scanning. Frame averages chosen for each image to yield SNR of approximately 35. (b) Line profile comparisons across different frame averaging for vessel shown at 750 μm depth in (a) for GG and RG scanning. Values of frame averaging shown in legend correspond to identical image acquisition times for each line color in legend across two the scanning methods. (c) Visual comparison of image quality at 750 μm depth across the two scanning methods for acquisition times of 3, 13.5 and 22.5 seconds.

For our imaging purposes, we determined that an SNR of 35 allows us to minimize acquisition time while preserving the abilities to effectively perform vectorization with our software (SLAVV) and visually discern all vessels for qualitative analysis.

### 3.3 Comparison with identical power levels and total dwell time

To provide a comparison between the resonant-galvo scanning and galvo-galvo scanning methods, a set of images were collected in which power at the sample and total sample exposure time were identical between the two methods at each depth of 250, 500, and 750 μm. The total sample exposure time is the sum of pixel dwell times across all pixels used to form the image. Table 1 compares the average SNR across ten vessels at each depth for sample exposure times of 1.89 and 5.66 seconds. Power levels at the sample were 5.8, 22, and 86 mW for 250, 500, and 750 μm depths, respectively. Paired t-test results showed that GG scanning produced higher SNR values at 500 μm depth with a sample exposure time of 1.89 seconds and at 750 μm with exposure times of 1.89 and 5.66 seconds (*p* < 0.05) compared to RG scanning. It should be noted that the mouse used for the measurements in Table 1 is not the same as that used for Figures 4 and 5.

**Table 1.**
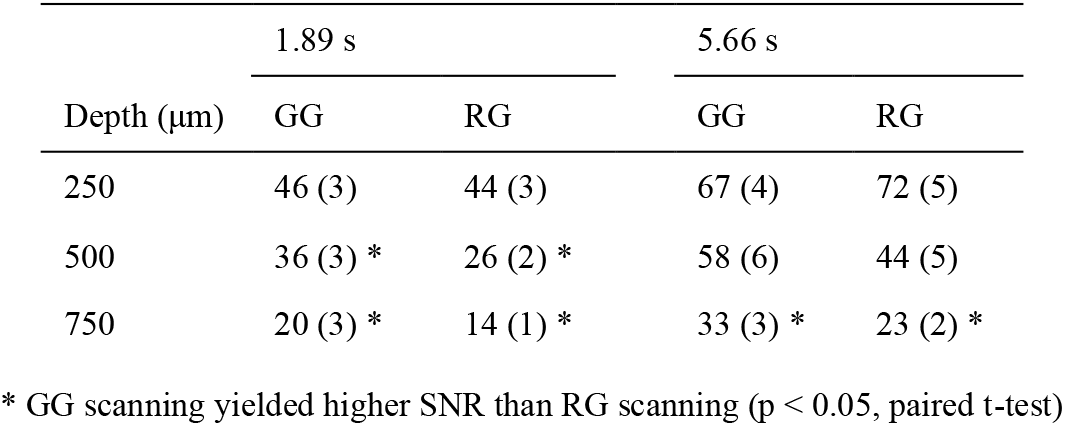
SNR Mean (Standard Error)

| Depth ($\mu\text{m}$) | 1.89 s | | 5.66 s | |
| --- | --- | --- | --- | --- |
|  | GG | RG | GG | RG |
| 250 | 46 (3) | 44 (3) | 67 (4) | 72 (5) |
| 500 | 36 (3) * | 26 (2) * | 58 (6) | 44 (5) |
| 750 | 20 (3) * | 14 (1) * | 33 (3) * | 23 (2) * |

\* GG scanning yielded higher SNR than RG scanning ( $p < 0.05$ , paired t-test)

### 3.4 Vectorization of two-photon vascular images

We used SLAVV software to examine whether the image signal to noise in RG image sets was sufficient for detailed analysis of vascular morphology. Figure 6(c) shows an example of a vectorized mosaic image (1050 μm × 1050 μm × 660 μm), which was formed by stitching together four 700 × 700 × 660 μm stacks using ImageJ [21]. Maximum intensity projections of this mosaic in the axial and sagittal directions are shown in Figures 6(a) and 6(b), respectively. The total acquisition time for this mosaic, collected using RG scanning, was 23 minutes (frame average of 20 was sufficient throughout for this particular mouse)—approximately a fourfold speedup compared to the expected GG acquisition time of 81 minutes for a stack of the same dimensions and image quality (assuming frame average of 2 throughout).

**Fig. 6.**
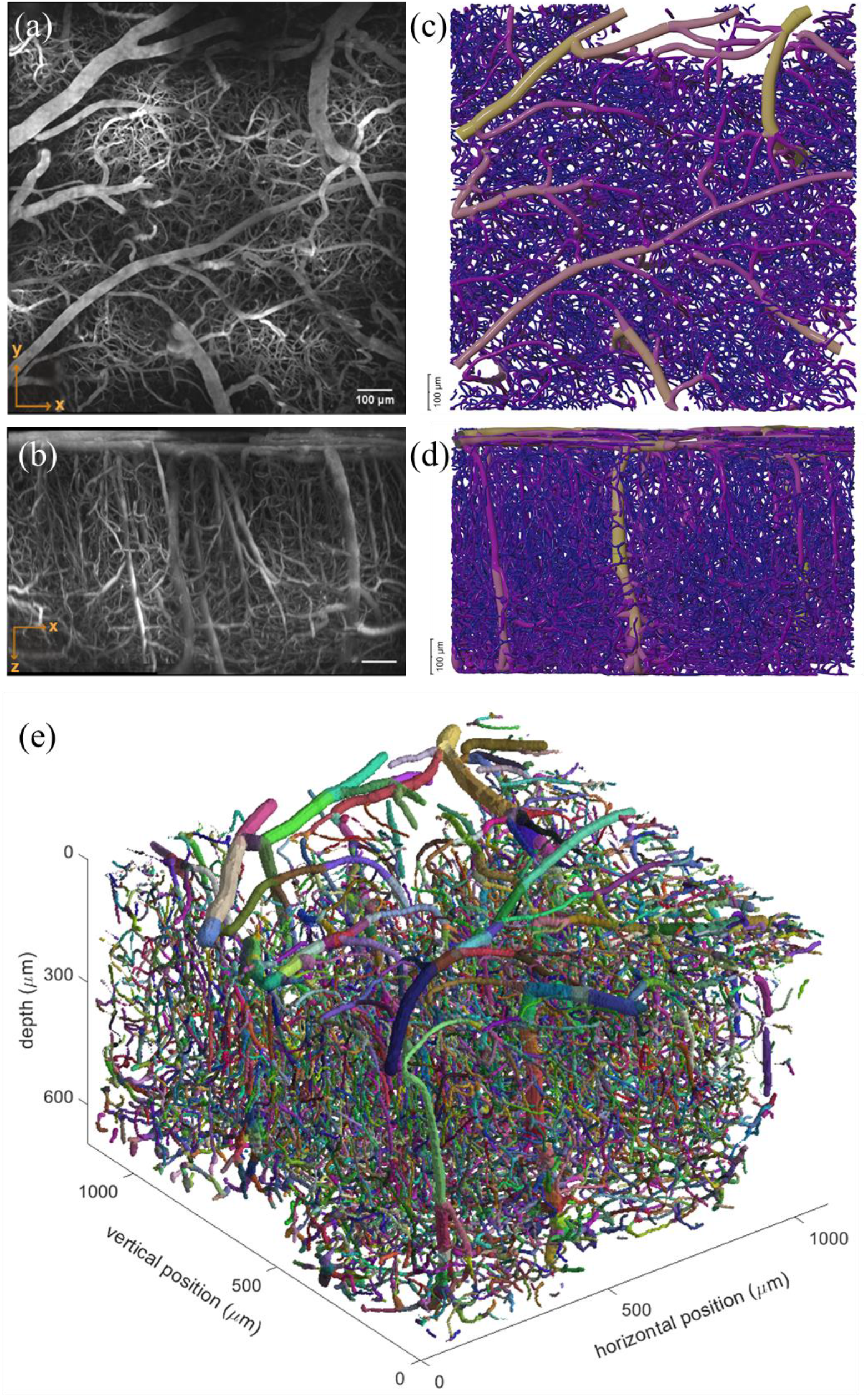
Large FOV mosaic imaging with resonant galvo acquired in 23 minutes. (a) maximum intensity x-y projection of tiled image (1050 μm × 1050 μm × 660 μm) comprised of four stacks of 700 μm × 700 μm × 660 μm stitched together using Fiji ImageJ. (b) maximum intensity x-z projection of image from (a). *VessMorphoVis* rendering of tiled image (color-coded by average section radius) after vectorization with SLAVV, represented as orthographic projections in (c) and (d), corresponding to projections (a) and (b). (e) Vectorized representation of vasculature in MATLAB from tiled image shown in (a-d) where each color denotes individually detected strands (defined as a segment between two branches).

## 4. Discussion

### 4.1 Signal-to-Noise Ratio Analysis

For a photomultiplier tube, the signal-to-noise ratio can be expressed as [25]:

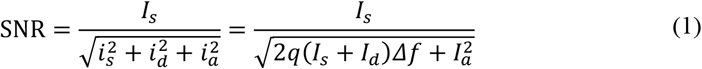

where the noise component is composed of photon shot noise (*i*_*s*_), dark current noise (*i*_*d*_), and transimpedance amplifier noise (*i*_*a*_). *I*_*s*_ represents the photocurrent created by light detection, *I*_*d*_ is the dark current, *I*_*a*_ is the input referred peak-to-peak noise current, *q* is the electron charge, and Δ*f* is the frequency bandwidth. *I*_*s*_ is directly related to the number of photons detected, which for two-photon imaging is proportional to the square of the average excitation power at the sample [25,26], assuming constant pulse duration and repetition rate. When the detection is shot-noise-limited, the SNR is thus linearly proportional to average excitation power [25,26]:

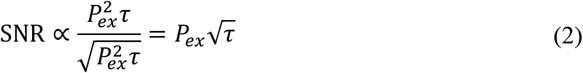

where (*P*_ex_) is the average excitation power at the sample in a time interval (*τ*).

The reduction in acquisition times using resonant-galvo scanning can be attributed to shot-noise-limited detection up to a certain depth, which allows for SNR to be maintained by using higher excitation powers with shorter pixel dwell times compared to galvo-galvo scanning (87.8 ns vs 7200 ns). To avoid tissue damage, these higher excitation powers cannot be used with GG scanning’s pixel dwell times, which are orders of magnitude higher than those of RG [16]. We observe a general linear relationship between SNR and excitation power at the sample in Figure 4 at depths of 250 μm and 500 μm, where a threefold increase in excitation power between GG and RG scanning at 250 μm resulted in a threefold increase in SNR and a twofold increase in power at 500 μm resulted in a twofold increase in SNR for identical imaging times (t = 3s). This data supports a shot-noise limited detection model as described in Eq. 2. At the depth of 750 μm, however, a 1.6x increase in power yields only a 1.5x increase in SNR. Given the weaker fluorescent signals at this greater imaging depth, it is likely that detection began to deviate from being shot-noise limited and for the SNR to be affected by other noise components: dark noise and amplifier noise as indicated in Eq. 1. A general linear relationship between SNR and the square root of acquisition time interval as described in Eq. 2 can also be seen in Figure 4 at depths of 250 μm and 500 μm—e.g., a fourfold increase in acquisition time, such as the acquisition of 2 frames vs. 8 frames for GG scanning or 20 frames vs. 80 frames for RG scanning, results in an approximate twofold increase in SNR as expected for shot-noise limited detection.

Results in Table 1 show that when using identical excitation power and total sample exposure times between the two scanning methods, GG scanning yields higher SNR values than RG scanning at deeper depths of 500 μm, with the shorter total sample exposure time of 1.89 seconds, and 750 μm, with both tested total sample exposure times of 1.89 and 5.66 seconds. Under presumed shot-noise-limited detection with sufficient signal strength at the 250 μm depth and with longer dwell time at the 500 μm depth, both methods yield statistically equivalent SNR values, as expected from Eq. 2. GG scanning outperforms RG scanning when detection is not shot-noise limited and is affected by the other noise components indicated in Eq. 1—dark noise and amplifier noise. The dark noise and amplifier noise are more significant in RG scanning, which requires a higher frequency bandwidth setting (14 MHz) and has greater input referred peak-to-peak noise (150 nA) compared to GG scanning (1 MHz, 2.4 nA). Thus, for situations in which excitation power for RG scanning cannot exceed what is used for GG scanning, such as for more sensitive samples, GG scanning can be expected to yield higher SNRs when detection is not shot-noise limited.

### 4.2 Difference in Temporal Resolution

Resonant-galvo scanning is an advantageous method for reducing acquisition times by up to sixfold per single image, compared to galvo-galvo scanning, while preserving image quality, provided that greater excitation power can be used without inducing tissue damage. With an RG frame rate of 28 Hz, each additional frame that we choose to average increases our acquisition time by only 36 milliseconds per axial slice, compared 2.1 seconds for each additional GG frame per slice. For each additional frame averaged in RG scanning, there is a small increase in SNR. With these small SNR increments, we can reach the exact minimum SNR needed for image analysis and thus minimize the acquisition time. GG scanning, in contrast, has a low temporal resolution that often forces us choose between under- or over-shooting the SNR because each additional frame averaged gives a larger SNR increase than with RG scanning. For instance, if the minimum required SNR for analysis is above what is achievable with two frames but under what can be achieved with three frames, we must spend the additional 2.1 seconds per axial slice to collect three frames to ensure that the SNR is at least above the minimum SNR required for analysis. In a typical acquisition of 251 axial slices for a total depth of 750 μm, the potential time spent overshooting SNR to prevent undershooting it could be several minutes for a GG stack, while no more than a few seconds for an RG stack. The high temporal resolution of RG scanning is especially advantageous in samples with low noise, where the number of frames needed to achieve the appropriate SNR requires only a small fraction of the time required by GG scanning to acquire just one frame. Thus, it should be noted that the exact time-savings of using RG scanning is dependent on sample quality, which can vary between animals, regions of interest, and even time of imaging.

### 4.3 Time Savings with Large FOV Imaging

The benefits of RG scanning are augmented when acquiring large field of view mosaic images, in which case the amount of time saved compounds. The single image stack acquired in Figure 3 using RG scanning was completed in 14 minutes. By interpolating the results from Figure 4, the amount of time that it would have taken GG scanning to collect the same image stack with the same minimum SNR is calculated to be 34 minutes. A significant 20 minutes were saved by using RG scanning for this single image. A large FOV image, however, is composed of multiple single images, and thus the time saved per single image is multiplied. For instance, the large FOV mosaic image in Figure 6 was collected in 23 minutes using RG scanning, in contrast to an expected 81 minutes required for a similar GG-acquired image. We saved an hour of experiment time and saved the mouse from an extra hour of anesthesia, of which long-term use has been shown to cause vascular dilation and could skew the collected data [27]. The shorter imaging time is further cost effective by reducing the amount of Texas Red dye needed for vascular imaging, since longer imaging sessions would require additional dye as it is cleared from the animal.

### 4.4 Microscope Design Advantages

A resonant-galvo-galvo system is valuable for allowing the seamless switch between GG, RG, and RGG scanning, each of which ideal for different applications [28]. A goal of our RGG design was to make the design adaptable to any custom system using available commercial parts. Our resonant scanner alignment method can be implemented with any resonant galvanometer and *xy*-galvanometer pair. Furthering accessibility is the relatively low cost of parts needed for the design. We calculate that the costs of the parts for our implementation of a resonant-galvo-galvo scanning system ($10,500) is approximately one-third the cost of purchasing a patented commercial scanner.

## 5. Conclusion

We demonstrate a simple-to-implement two-photon microscope design that allows effortless switching between galvo-galvo and resonant-galvo scanning. We compared the performance between GG and RG scanning and found that the SNR for both scan modes behave as expected for a system with shot-noise limited detection when sufficient fluorescent signal is present. We show that RG scanning can reduce the acquisition time from that of GG scanning for a desired SNR when the excitation power can be increased without causing tissue damage. The limitations of this acquisition speedup will vary with a variety of factors, such as imaging depth, and are expected to be sample-dependent. When excitation power must be maintained at levels used for GG scanning, GG scanning performs similarly to RG scanning at depths where detection is shot-noise limited but outperforms RG scanning in achieving desired SNR in less acquisition time at deeper imaging depths where detection deviates from being shot-noise limited.

## Acknowledgments

We thank Thorlabs for granting permission to reproduce the schematic of the galvo-galvo pair in Figure 2. We also thank Dr. Marwan Abdellah for help and guidance with *VessMorphoVis* software.

## Data availability

Data underlying the results presented in this paper are not publicly available at this time but may be obtained from the authors upon reasonable request.

## Supplemental document

See Supplement 1 for supporting content.

## References

1. F. Helmchen and W. Denk, “Deep tissue two-photon microscopy,” Nat Methods 2, 932–940 (2005).

2. Hanley, Verveer, Gemkow, Arndt-Jovin, and Jovin, “An optical sectioning programmable array microscope implemented with a digital micromirror device,” J Microsc 196, 317–331 (1999).

3. V. Bansal and P. Saggau, “Digital Micromirror Devices: Principles and Applications in Imaging,” Cold Spring Harbor Protocols 2013, 404–411 (2013).

4. G. Duemani Reddy, K. Kelleher, R. Fink, and P. Saggau, “Three-dimensional random access multiphoton microscopy for functional imaging of neuronal activity,” Nature Neuroscience 11, 713–720 (2008).

5. R. Y. Tsien and B. J. Bacskai, “Video-Rate Confocal Microscopy,” in Handbook of Biological Confocal Microscopy, J. B. Pawley, ed. (Springer US, 1995), pp. 459–478.

6. M. B. Bouchard, V. Voleti, C. S. Mendes, C. Lacefield, W. B. Grueber, R. S. Mann, R. M. Bruno, and E. M. C. Hillman, “Swept confocally-aligned planar excitation (SCAPE) microscopy for high-speed volumetric imaging of behaving organisms,” Nature Photon 9, 113–119 (2015).

7. V. Voleti, K. B. Patel, W. Li, C. Perez Campos, S. Bharadwaj, H. Yu, C. Ford, M. J. Casper, R. W. Yan, W. Liang, C. Wen, K. D. Kimura, K. L. Targoff, and E. M. C. Hillman, “Real-time volumetric microscopy of in vivo dynamics and large-scale samples with SCAPE 2.0,” Nat Methods 16, 1054–1062 (2019).

8. D. R. Beaulieu, I. G. Davison, K. Kılıç, T. G. Bifano, and J. Mertz, “Simultaneous multiplane imaging with reverberation two-photon microscopy,” Nat Methods 17, 283–286 (2020).

9. J. L. Fan, J. A. Rivera, W. Sun, J. Peterson, H. Haeberle, S. Rubin, and N. Ji, “High-speed volumetric two-photon fluorescence imaging of neurovascular dynamics,” Nat Commun 11, 6020 (2020).

10. J. Wu, Y. Liang, S. Chen, C.-L. Hsu, M. Chavarha, S. W. Evans, D. Shi, M. Z. Lin, K. K. Tsia, and N. Ji, “Kilohertz two-photon fluorescence microscopy imaging of neural activity in vivo,” Nat Methods 17, 287–290 (2020).

11. S. Xiao, I. Davison, and J. Mertz, “Scan multiplier unit for ultrafast laser scanning beyond the inertia limit,” Optica 8, 1403 (2021).

12. P. Mahou, J. Vermot, E. Beaurepaire, and W. Supatto, “Multicolor two-photon light-sheet microscopy,” Nat Methods 11, 600–601 (2014).

13. T. A. Pologruto, B. L. Sabatini, and K. Svoboda, “ScanImage: Flexible software for operating laser scanning microscopes,” BioMed Eng OnLine 2, 13 (2003).

14. S. A. Engelmann, A. Zhou, A. M. Hassan, M. R. Williamson, J. W. Jarrett, E. P. Perillo, D. J. Spence, T. A. Jones, and A. K. Dunn, “Diamond Raman Laser and Yb Fiber Amplifier for In Vivo Multiphoton Fluorescence Microscopy,” bioRxiv (2021).

15. E. P. Perillo, J. W. Jarrett, Y.-L. Liu, A. Hassan, D. C. Fernée, J. R. Goldak, A. Bonteanu, D. J. Spence, H.-C. Yeh, and A. K. Dunn, “Two-color multiphoton in vivo imaging with a femtosecond diamond Raman laser,” Light Sci Appl 6, e17095–e17095 (2017).

16. K. Podgorski and G. Ranganathan, “Brain heating induced by near-infrared lasers during multiphoton microscopy,” Journal of Neurophysiology 116, 1012–1023 (2016).

17. M. Desjardins, K. Kılıç, M. Thunemann, C. Mateo, D. Holland, C. G. L. Ferri, J. A. Cremonesi, B. Li, Q. Cheng, K. L. Weldy, P. A. Saisan, D. Kleinfeld, T. Komiyama, T. T. Liu, R. Bussell, E. C. Wong, M. Scadeng, A. K. Dunn, D. A. Boas, S. Sakadžić, J. B. Mandeville, R. B. Buxton, A. M. Dale, and A. Devor, “Awake Mouse Imaging: From Two-Photon Microscopy to Blood Oxygen Level–Dependent Functional Magnetic Resonance Imaging,” Biological Psychiatry: Cognitive Neuroscience and Neuroimaging 4, 533–542 (2019).

18. D. R. Miller, A. M. Hassan, J. W. Jarrett, F. A. Medina, E. P. Perillo, K. Hagan, S. M. Shams Kazmi, T. A. Clark, C. T. Sullender, T. A. Jones, B. V. Zemelman, and A. K. Dunn, “In vivo multiphoton imaging of a diverse array of fluorophores to investigate deep neurovascular structure,” Biomedical Optics Express 8, 3470 (2017).

19. E. P. Perillo, J. E. McCracken, D. C. Fernée, J. R. Goldak, F. A. Medina, D. R. Miller, H.-C. Yeh, and A. K. Dunn, “Deep in vivo two-photon microscopy with a low cost custom built mode-locked 1060 nm fiber laser,” Biomed. Opt. Express 7, 324–334 (2016).

20. C. J. Schrandt, S. S. Kazmi, T. A. Jones, and A. K. Dunn, “Chronic Monitoring of Vascular Progression after Ischemic Stroke Using Multiexposure Speckle Imaging and Two-Photon Fluorescence Microscopy,” J Cereb Blood Flow Metab 35, 933–942 (2015).

21. J. Schindelin, I. Arganda-Carreras, E. Frise, V. Kaynig, M. Longair, T. Pietzsch, S. Preibisch, C. Rueden, S. Saalfeld, B. Schmid, J.-Y. Tinevez, D. J. White, V. Hartenstein, K. Eliceiri, P. Tomancak, and A. Cardona, “Fiji: an open-source platform for biological-image analysis,” Nat Methods 9, 676–682 (2012).

22. J. Ahrens, B. Geveci, and C. Law, “ParaView: An End-User Tool for Large Data Visualization,” in Visualization Handbook, C. D. Hansen and C. R. Johnson, eds. (Butterworth Heinemann, 2005), pp. 717–731.

23. S. A. Mihelic, W. A. Sikora, A. M. Hassan, M. R. Williamson, T. A. Jones, and A. K. Dunn, “Segmentation-Less, Automated, Vascular Vectorization,” PLoS Comput Biol 17, e1009451 (2021).

24. M. Abdellah, N. R. Guerrero, S. Lapere, J. S. Coggan, D. Keller, B. Coste, S. Dagar, J.-D. Courcol, H. Markram, and F. Schürmann, “Interactive visualization and analysis of morphological skeletons of brain vasculature networks with VessMorphoVis,” Bioinformatics 36, i534–i541 (2020).

25. E. L. Dereniak and D. G. Crowe, Optical Radiation Detectors, Wiley Series in Pure & Applied Optics (Wiley, 1984).

26. C. Xu and W. W. Webb, “Measurement of two-photon excitation cross sections of molecular fluorophores with data from 690 to 1050 nm,” J. Opt. Soc. Am. B 13, 481 (1996).

27. R. Cao, J. Li, B. Ning, N. Sun, T. Wang, Z. Zuo, and S. Hu, “Functional and oxygen-metabolic photoacoustic microscopy of the awake mouse brain,” NeuroImage 150, 77–87 (2017).

28. C.-H. Yu, J. N. Stirman, Y. Yu, R. Hira, and S. L. Smith, “Diesel2p mesoscope with dual independent scan engines for flexible capture of dynamics in distributed neural circuitry,” bioRxiv 2020.09.20.305508 (2020).

